# fastdemux: Robust SNP-based demultiplexing of single-cell population genomics data

**DOI:** 10.64898/2026.02.10.705082

**Authors:** Ali Ranjbaran, Francesca Luca, Roger Pique-Regi

## Abstract

Sample multiplexing reduces cost and batch effects in population based large-scale single-cell genomics studies but requires accurate and scalable computational demultiplexing. Existing genotype-based methods, such as demuxlet, provide high accuracy but can be computationally slow and memory intensive as the number of cells, donors, and informative variants increases. Here, we introduce `fastdemux,` a scalable genotype-based demultiplexing framework based on a diagonal linear discriminant analysis (DLDA) model that substantially improves computational efficiency while maintaining accurate donor assignment. Using a pooled single-cell RNA-seq dataset from unrelated donors, we benchmarked `fastdemux` against demuxlet, vireo, and demuxalot.`fastdemux` achieved comparable or improved demultiplexing accuracy while reducing runtime and peak memory usage by orders of magnitude relative to alternative methods. Performance remained robust across varying sequencing depths and genotype SNP filtering thresholds. In addition, the DLDA framework naturally extends to doublet and higher-order multiplet detection. We also show that`fastdemux` works well with scATAC-seq data where genetic variants are more sparsely covered. Together, these results establish `fastdemux` as an efficient and scalable solution for genetic demultiplexing of pooled single-cell datasets.

## Introduction

Over the past decade, advances in single-cell genomics have transformed our ability to characterize complex biological systems with high resolution. Traditional bulk sequencing assays measure average signals across millions of cells in a given tissue, masking the substantial cellular heterogeneity that underlies tissue function, developmental trajectories, and disease processes. In contrast, single-cell technologies such as single-cell RNA sequencing (scRNAseq), single-cell ATAC sequencing (scATAC-seq), and emerging multi-omic assays capture molecular profiles at the level of individual cells [9, 18, 14]. Single-cell technologies, therefore, enabled the discovery of rare subpopulations, cell-state transitions, and regulatory programs that are indistinguishable in bulk measurements. scRNA-seq has become essential for diverse applications, including mapping cell-type diversity across human tissues and building large-scale reference atlases [4, 34, 28], dissecting tumor microenvironments [13], characterizing dynamic responses to genetic or pharmacological perturbations at scale [2, 6], and enabling population-scale studies that integrate cellular phenotypes with genetic variation, including single-cell eQTL mapping and related frameworks [36, 5, 31]. As single-cell assays become increasingly high-throughput and cost-efficient, the scale at which experiments are performed, ranging from large tissue atlases to population-level cohorts, continues to expand, underscoring the need for robust computational methods to extract reliable biological insights from these massive and heterogeneous datasets [8, 18].

Despite the rapid adoption of single-cell technologies, several experimental and technical challenges continue to limit their scalability and interpretability, particularly in large cohort and population-based studies. Single-cell experiments remain resource-intensive, as library preparation reagents, sample handling, and sequencing requirements scale with the number of samples and experimental batches, constraining study size even as per-base sequencing costs decline. These challenges are compounded by intrinsic features of single-cell data, including sparsity and dropout, which increase technical noise and reduce sensitivity for detecting subtle biological signals [24, 39, 32, 21, 23]. Collectively, these limitations motivate the development of experimental and computational strategies that reduce cost, reduce technical variability, and improve comparability across samples.

To address the cost and batch-related challenges of large single-cell studies, sample multiplexing has emerged as a widely adopted experimental strategy. In multiplexed designs, cells from multiple biological samples or individuals are combined and processed within one single-cell library preparation, thereby reducing per-sample reagent while minimizing technical variability introduced across batches. Post-sequencing demultiplexing is then required for computational assignment of individual cells back to their sample of origin. The first widely adopted framework for demultiplexing was introduced by demuxlet, which leverages naturally occurring genetic variation to infer donor identity directly from single-cell RNA-seq data [11]. Demuxlet assigns each cell captured in a droplet to its individual source by computing the maximum likelihood of observing RNA-seq reads overlapping known single-nucleotide polymorphisms (SNPs), given the reference genotypes of each donor, thereby enabling both donor identification and detection of doublets (droplets with two or more cells). Detecting doublets is easier when multiplexing a large number of donors, as the probability of doublets formed from cells from different individuals is much higher than those that come from the same individual [11]. Therefore, multiplexing allows to load a much higher number of cells in droplet based systems while still being able to accurately detect doublets. This combinatorial advantage is what allows loading libraries with a large number of cells.

Multiplexing study designs that utilize genetic variation for donor assignments can be particularly useful for population level studies to significantly reduce cost. Such genotype-based demultiplexing approaches have been applied in prior studies of human tissues, including placenta samples in human pregnancy research [35, 27] where fetal and maternal cells can then be discriminated by these approaches. These study designs have also been implemented to study psychosocial environments across immune cell populations in large cohorts [3]. Demultiplexing by genotype can be used in large/case control cohorts to make single-cells studies more feasible for clinical and epidemiological applications [12, 37, 30, 25, 7].

In addition to demultiplexing approaches that rely on available genotype reference data, a complementary class of genotype-free demultiplexing methods has been developed to enable sample assignment when donor genotypes are unavailable. These methods infer donor identities directly from allele frequency patterns observed in pooled single-cell data, typically using Bayesian probabilistic clustering, maximum likelihood, or matrix factorization-based frameworks [17, 10, 38]. However, scRNA-seq data contain sparse and uneven coverage, which makes accurate variant identification more challenging in the absence of genotype priors and can reduce performance under noisy or complex conditions [15, 26]. Beyond the use of genetic variation for demultiplexing, experimental strategies have also been developed that rely on nucleotide-barcoded antibodies [22, 20] or membrane-anchored oligonucleotides [29, 16] to label cells or nuclei prior to pooling. However these experimental approaches are less widely used and are less versatile, for example they are not compatible with some of the scATAC-seq or single nuclei protocols, may require fixing or sorting which can introduce an unwanted perturbation, and label leaching may also occur. These methods are still useful in situations where the same donor is used (identical genotypes) but a large set of conditions/treatments is explored. Nevertheless, here we will focus on leveraging genetic variation for donor multiplexing, which can be applied to a wide range of single-cell protocols and on population based studies where a large number of donors are available or desirable.

The most widely used demultiplexing tools such as demuxlet, face important computational limitations as single-cell studies continue to scale in size. When applied to a large number of cells and donors, they are computationally slow and memory-intensive. In addition, current tools’ performance in high-throughput computing environments is further constrained by the limited support for parallel execution in multiple CPUs. Many of the existing tools are also not readily compatible with scATACseq, and when they are, due to the nature of the data, the computational issues are exacerbated as more genetic variants across the genome (not only in the transcriptome) need to be studied with only one or two fragments covering them per cell barcode. To address these challenges, we introduce `fastdemux,` a scalable demultiplexing framework based on a diagonal linear discriminant analysis (DLDA) model that substantially reduces computational overhead and memory requirements while preserving accurate donor assignment. The DLDA score calculation is implemented as a streaming algorithm, meaning that it is calculated as the files with the genotypes and read alignments are being read and therefore do not require multiple iterations or loading large amounts of data beforehand. Our benchmarking analysis shows that by improving runtime and memory efficiency,`fastdemux` enables genetic demultiplexing to be applied more broadly to large pooled single-cell datasets compared to existing methods.

## Methods

### fastdemux model

`fastdemux` is a genotype-based single-cell demultiplexing method based on a diagonal linear discriminant analysis (DLDA) framework. The basic idea is that for each barcode we collect all the reads overlapping genetic variants, and we build a discriminant function for each barcode/individual pair. Generally, we will assign the barcode to the individual with the highest scoring discriminant function (except the case where a doublet or higher-order multiplet discriminant gives a higher score as it will be detailed later).

The method takes as input a genotype VCF file (with *L* biallelic variants and *N* individuals), an aligned BAM file produced by the 10x Genomics Cell Ranger pipeline, and a set of cellular barcodes. This can be the full unfiltered list of barcodes or a filtered list of *B* barcodes.

Let *b* = 1,. .., *B* index cell barcodes and *n* = 1,. .., *N* index individuals (donors). For each barcode *b*, reads overlapping genetic variants are indexed by variant (biallelic SNPs) *l* = 1,. .., *L* and read *k* = 1,. .., *K*_*bl*_, where *K*_*bl*_ denotes the number of reads from barcode *b* overlapping variant *l*.

Each read-level observation for the BAM file *X*_*bk*_ is treated as independent and encoded as

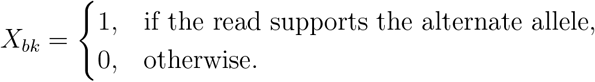

For each individual *n* and variant *l*, the genotype dosage is denoted by *d*_*nl*_ ∈ {0, 1, 2 } The alternate allele frequency at variant *l* is defined as

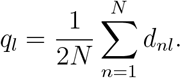

Note that the allelic dosage can be defined as floating point numbers to account for genotype uncertainty and is usually defined in the DS field of the vcf file by imputation and genotyping methods. If DS is not defined in the vcf file we will use genotype probabilities (GP field) or the vcf GT field if GP is not present. SNPs are only considered if they have at least one read covering them in the single cell dataset, and genotype information is available for at least two individuals in the vcf file. Individuals with a missing genotype or not valid DS/GP/GT fields will be imputed to the overall mean dosage *d*_*nl*_ = 2*q*_*l*_. However, it is always possible to further filter the vcf file requiring higher coverage or removing SNPs with missing genotypes. Currently only biallelic SNPs are taken into account and any other type of genetic variant in the vcf file will be ignored.

Then for each barcode *b* and individual *n*, the DLDA discriminant score is defined as

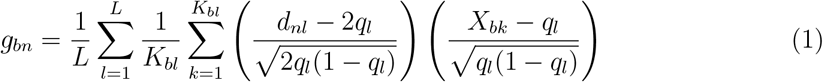

Each read *X*_*bk*_ is treated as conditionally independent given the barcode, and both the genotype-based weight and the read-level observation are normalized to have mean zero and variance one (assuming that would be randomly sampled from any individual in the vcf file). Note that the left parentheses with the genotype dosage is exactly the same that is typically used to calculate the principal components (PCs) on the genotype data. Compared to the calculation of PCs where we correlate centered genotype data from each pair of individuals, in DLDA we correlate the genotype derived from the reads in each barcode, to the genotype in the vcf file for each individual. This can also be seen more readily after aggregating reads at each variant,

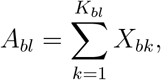

as the singlet DLDA score can be equivalently written as

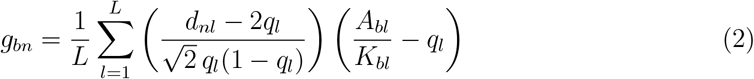

Calculating these discriminants is also very computationally and memory efficient. Assuming the BAM file and vcf files are ordered consistently, the summations in the discriminant can be calculated while streaming over the two files, so very minimal memory is necessary scaling linearly with the number of barcodes and individuals in the vcf file 𝒪 (*NB*). Computation also scales linearly with the number of reads overlapping genetic variants, that is with the size of the BAM file and the number of rows in the vcf file, more specifically with the number of genetic variants with reads covering them 𝒪 (*L*).

### DLDA for doublets and higher-order multiplets

Genotype information can be used to identify droplets that contain more than one cell when they are from different donors. Using the DLDA framework we can calculate the correlation we would get between the reads, if the barcode can randomly sample genotypes from two individuals. For a barcode *b* arising from a doublet of individuals *n* and *n*′, the DLDA score is defined as

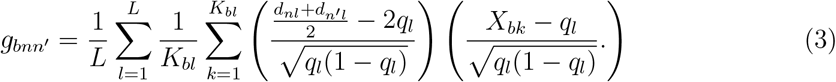

After aggregating reads at each variant, this becomes

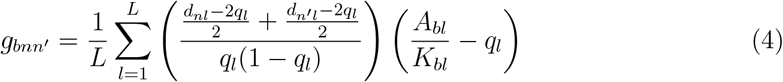

The expected dosage would be the average of the two individual dosages and the variance of the centered dosage is half that of the singlet. The centered read dosage would be exactly the same as for a singlet. Importantly, the doublet discriminant score can be expressed directly in terms of the corresponding singlet discriminant scores as:

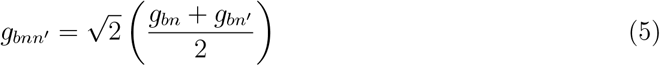

`fastdemux` naturally generalizes to higher-order multiplets (M-lets). For a barcode *b* composed of *M* individuals *n*_1_,. .., *n*_*M*_, the DLDA discriminant score is defined as:

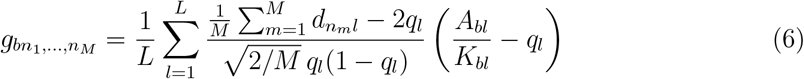

Equivalently, the M-let score can be expressed directly in terms of singlet scores as

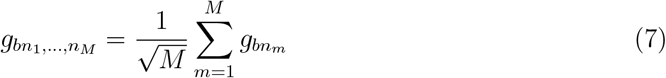

This formulation allows`fastdemux` to evaluate singlet, doublet, and higher-order multiplet hypotheses within a unified DLDA framework while maintaining computational efficiency. Naturally the best scoring doublet would consist of the individuals in the top 2 scoring singlets, and the best scoring *M* -let would be of the top *M* singlets. Thus, we can calculate if an *M* -let or a singlet provides a better discriminant at the end, if we record all the discriminants, requiring 𝒪 (*NB*) memory and computation.

### Benchmarking dataset

To evaluate the demultiplexing performance of `fastdemux` relative to other genotype-based demultiplexing tools, we analyzed a pooled single-cell RNA-seq dataset derived from unrelated Yoruba individuals from Ibadan, Nigeria (YRI) [19]. This pool was generated from heterogeneous differentiating cultures (HDCs), an in vitro system in which pluripotent cells asynchronously differentiate into a broad spectrum of cell types. We selected a single pool containing eight donors (five females and three males), corresponding to the largest donor pool available in the study.

Raw sequencing data (FASTQ files) were downloaded from the Sequence Read Archive (SRA) using sample 1019 (experiment SRX24003763; BioSample SAMN40555532; SRA study SRS20800172). This dataset was previously described in [19] and provides a well-characterized benchmark for evaluating both demultiplexing accuracy and computational performance in pooled single-cell experiments. All FASTQ files were processed uniformly across demultiplexing tools to ensure comparability. A total of 1,057,638,835 reads from this library were aligned to the GRCh38.p12 human reference genome using Cell Ranger, resulting in 13,873 estimated cells from 8 donors. Cells with fewer than 500 features were filtered out for downstream analysis.

Genotype data for the corresponding donors were obtained from the 1000 Genomes Project [1]. The genotype files used in this assessment included all 53 unrelated individuals from this population. For the baseline analysis, genotypes were filtered to include only biallelic SNPs with at least 10 supporting reads. Demultiplexing was performed using demuxlet 1.0, vireo 0.5.8, demuxalot 0.4.1, and `fastdemux` 0.1.1 with corresponding default parameters. For consistency across all demultiplexing runs, we allocated 2 CPUs and 510 GB of memory, but all methods were run on a single thread setting.

We also evaluated the method with scATAC-seq data for a multiplexed library of LPS-treated PBMC from 12 individuals [3]. We used `fastdemux` to demultiplex this overloaded library of 20,000 cells with 1,298,161,016 raw fragments, 64,908.05 mean fragments per cell, 22,948 median high-quality fragments per cell, and 94.0% fraction of high-quality fragments in cells.

### Expression-based sex inference and misclassification analysis

To assess agreement between donor sex metadata and expression-based sex inference, we used raw RNA count matrices to compute per-cell X/Y expression ratio by summing counts across genes located on the X and Y chromosomes, with a pseudocount added prior to log_10_ transformation. Unsupervised k-means clustering (*k* = 2) was applied to the log-transformed ratios to identify male- and female-like expression clusters, and the midpoint between cluster centroids was used as a global threshold for sex assignment. This threshold was computed once using the pooled dataset and applied uniformly across all demultiplexing tools. For each tool, expression-predicted sex was compared to donor sex metadata, and cells lacking either annotation were excluded. Sex misclassification was defined as disagreement between expression-predicted sex and donor sex metadata, and misclassification rates were calculated as the fraction of misassigned cells per tool.

### Read subsampling

To assess demultiplexing robustness under reduced sequencing depth, we performed a read subsampling analysis using fractions of the original BAM files (1,057,638,835 total reads and 76,237 mean reads per cell) corresponding to 1% (10,576,388), 5% (52,881,941), 10% (105,763,883), 30% (317,291,650), and 50% (528,819,417) of reads. For each fraction, sub-sampled BAM files were generated and processed independently through demuxlet, vireo, demuxalot, and `fastdemux` using identical parameters to the full-depth analysis. Demultiplexing outputs were merged into Seurat objects, and donor assignments were extracted for each tool and fraction. A consensus donor label was constructed using full-depth (100%) data by applying a union-based agreement strategy across tools, requiring agreement among multiple tools when available. This consensus served as a reference for evaluating fraction-level assignments. Performance metrics were computed per tool and fraction by comparing fraction-level calls to the consensus donor labels.

### VCF SNP-filtering analysis

To evaluate sensitivity to genotype reference quality, we performed demultiplexing using genotype VCF files filtered at progressively more stringent SNP thresholds (G199, G99, G49, and G9), excluding unfiltered (G0) variants. For each demultiplexing tool, analyses were run independently at each SNP threshold. Tool-specific assignments generated using the G9-filtered VCF were used as an internal baseline reference for that tool. For each SNP threshold, donor assignments were compared to the corresponding G9 assignments on a percell basis, restricting comparisons to droplets assigned in both conditions. Assignment rates were computed per tool and filter level, and the total-droplet error rate was calculated as one minus the fraction of correctly assigned droplets relative to all droplets, enabling direct comparison across tools and filtering conditions.

## Results

We benchmarked the computational performance of`fastdemux` against other genotype-based demultiplexing tools, including demuxlet, vireo, and demuxalot, using an 8-donor pooled scRNA-seq dataset of HDCs of unrelated Yoruba individuals from Ibadan, Nigeria (YRI) [19]. Genotype data for these individuals were also available through the 1000 Genomes Project [1]. Total elapsed runtime and peak memory usage were recorded for each tool under comparable computational settings (Fig. 1A–B; Table 1).`fastdemux` completed demultiplexing substantially faster than commonly used methods, with a total elapsed run-time of approximately 42 hours, compared to ∼ 210 hours for demuxlet and∼ 302 hours for vireo. Demuxalot exhibited the shortest runtime overall (∼ 15 hours), but required substantially higher memory than `fastdemux.` In terms of memory usage, `fastdemux` showed a marked reduction in peak resident memory, with a maximum resident set size (RSS) of ∼ 1 GB, compared to ∼ 49 GB for demuxalot and *>*460 GB for demuxlet and vireo (Fig. 1B). These results demonstrate that `fastdemux` achieves favorable scaling in both runtime and memory consumption relative to existing approaches.

**Figure 1.**
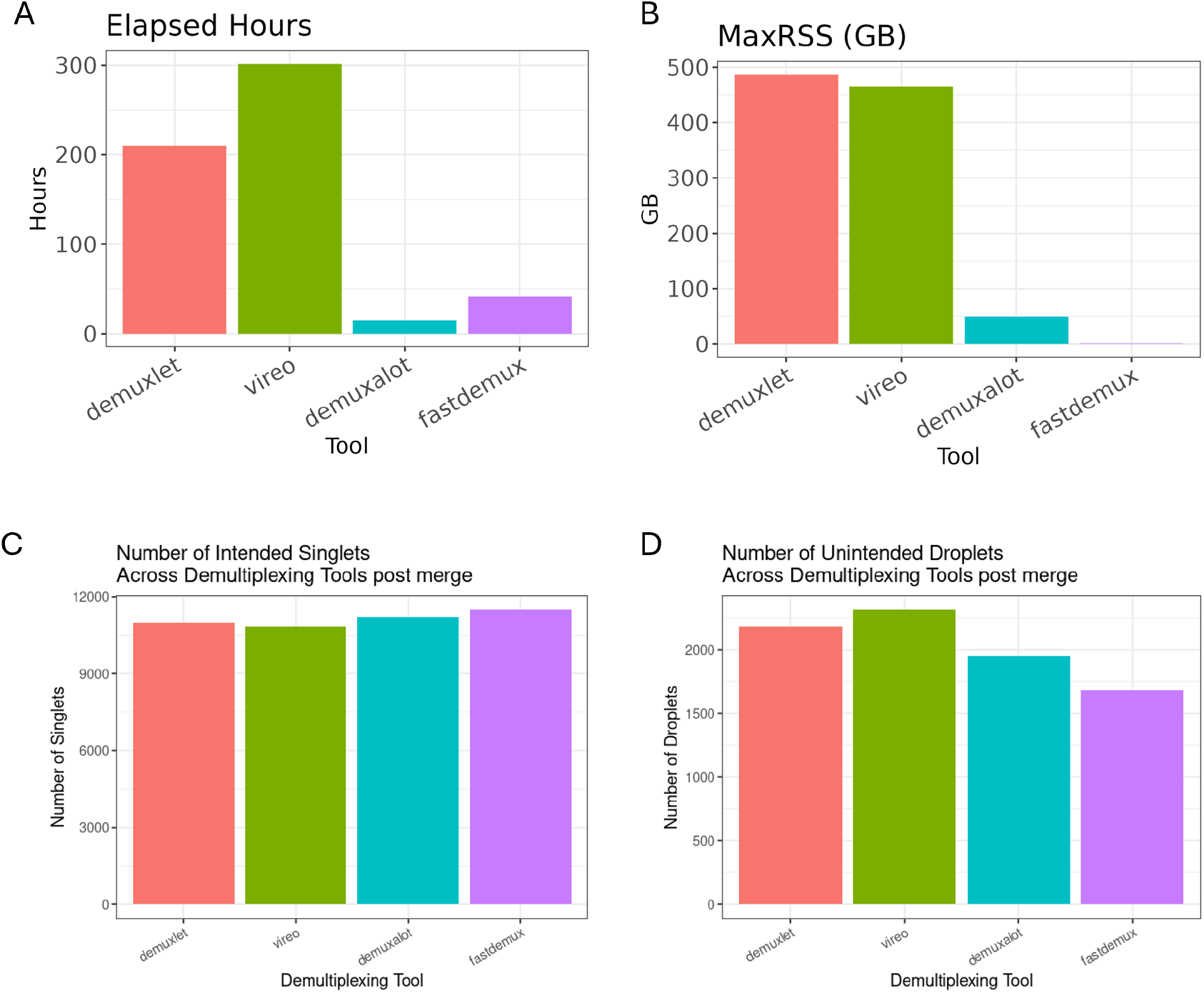
Computational performance and demultiplexing outcomes across tools in an 8-donor pooled scRNA-seq dataset. (A) Total elapsed runtime (hours) and (B) peak resident memory usage (MaxRSS, GB) for demuxlet, vireo, demuxalot, and `fastdemux` under comparable computational settings. (C) Number of intended singlets recovered and (D) number of unintended droplets (doublets and higher-order multiplets) assigned by each method after merging demultiplexing outputs. All analyses were performed on the same pooled 8-donor scRNA-seq dataset.

We next evaluated demultiplexing outcomes across tools by comparing the number of intended singlets and unintended droplets assigned (Fig. 1C–D; Tables 2–3). All tools recovered a comparable number of intended singlets, with `fastdemux` identifying the largest number of intended singlets (11,490), slightly exceeding demuxlet (10,987), vireo (10,858), and demuxalot (11,219). The fraction of intended singlets among assignments was highest for `fastdemux` (0.87), compared to 0.83 for demuxlet, 0.82 for vireo, and 0.85 for demuxalot (Table 3). Correspondingly, `fastdemux` assigned the lowest fraction of unintended droplets (0.13). Donor-level recovery was highly consistent across tools, with similar numbers of singlets recovered per individual donor (Supplementary Fig. S1; Table 4), indicating that differences in computational performance were not accompanied by systematic biases in donor assignment. Together, these results demonstrate that`fastdemux`maintains demultiplexing accuracy comparable to or exceeding existing widely used methods while substantially reducing computational resource requirements.

To evaluate concordance between donor sex metadata and expression-based sex inference, we constructed a global expression-derived sex reference using the pooled scRNA-seq dataset. Sex inference was derived independently of demultiplexing using a global expression-based X/Y ratio reference. Expression-based sex inference yielded similar misclassification rates across all demultiplexing tools (Fig. S2). `fastdemux` exhibited a sex misclassification rate comparable to demuxlet, vireo, and demuxalot, with no substantial reduction in misclassification relative to existing methods. Differences in misclassification rates across tools were modest and consistent with variability expected from expression-based sex inference in scRNA-seq data. These results indicate that sex misclassification is primarily driven by limitations inherent to expression-level sex inference, such as low Y-chromosome expression, rather than differences in demultiplexing performance. Importantly,`fastdemux` does not introduce systematic sex assignment bias relative to other methods.

To evaluate how demultiplexing performance depends on sequencing depth, we compared all tools using downsampled fractions (1%, 5%, 10%, 30%, and 50%) of the original reads (1,057,638,835). Across all tools, demultiplexing accuracy improved with increasing read fraction, with the highest error rates observed at the lowest sequencing depths for all tools (1– 5%) (Fig. 2A). At very low read fractions, `fastdemux` did not outperform other methods and exhibited error rates comparable to demuxlet, vireo, and demuxalot. However, at higher read fractions (≥30%), `fastdemux` consistently achieved lower error rates than the other tools. At 30% and 50% of reads, `fastdemux` showed the largest reduction in total-droplet error rate, indicating improved robustness as sequencing depth increased. `fastdemux` completed demultiplexing using all tested read fractions substantially faster than demuxlet and vireo (Fig. 2B, Table 5). `fastdemux` also had a significant marked reduction in peak resident memory, with an RSS of less than 1GB across all read fractions, compared to∼22 GB for demuxalot, *>*42 GB for demuxlet, and *>* 27GB for vireo even at 1% read fraction (Fig. 2C, Table 5). These results demonstrate that while all methods are sensitive to extreme read downsampling,`fastdemux` more effiectively leverages larger sequencing depth to achieve accurate donor assignment.

**Figure 2.**
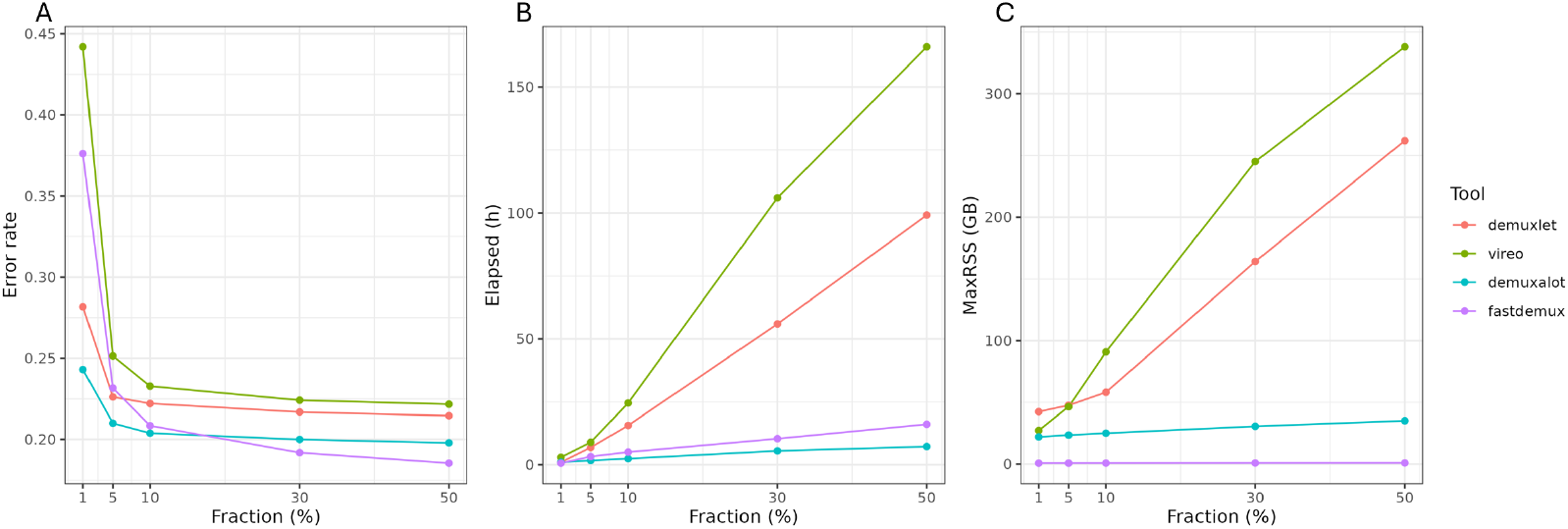
Demultiplexing performance as a function of sequencing depth. x-axis represents the fraction of the total reads downsampled, and the y-axis compares across demultiplexing tools: (A) Total-droplet error rate, defined as 1 − (correctly assigned droplets*/*total droplets). (B) Total elapsed runtime in hours. (C) Peak resident memory usage in gigabytes (MaxRSS). All tools were evaluated on the same pooled 8-donor scRNA-seq dataset.

In practice, when considering biallelic genetic variants for demultiplexing, high-coverage SNPs are often preferred because they tend to be more informative for donor assignment. Restricting analyses to higher-coverage SNPs (fewer SNPs) also significantly reduces runtime and memory usage by limiting the number of variants evaluated; however, even a larger number of SNPs at lower coverage can still be very informative in discriminating the best matching individual. To determine how reductions in the number of SNPs considered affect demultiplexing accuracy, we filtered the genotype file at multiple SNP coverage thresholds and performed demultiplexing for each filtered genotype input. All demultiplexing tools exhibited increasing error rates as the number of retained SNPs decreased, reflecting reduced genotype information available for donor assignment (Fig. 3A). Across all filtering thresholds, `fastdemux` consistently showed lower total-droplet error rates relative to demuxlet, vireo, and demuxalot when compared against each tool’s default run at greater than nine SNPs of coverage (G9). While performance differences were modest at higher SNP coverage thresholds (G199 and G99), all tools exhibited a similar trend of decreasing error rates as more SNPs are used (i.e. when adding SNPs with lower coverage). Differences between methods became more apparent at lower SNP thresholds (G49 and G9), with `fastdemux` retaining the lowest error rate. Overall, these findings indicate that `fastdemux` is comparatively robust to reductions in genotype SNP density. More importantly, we demonstrate that `fastdemux` permits the inclusion of a large number of lower-coverage SNPs that remain informative for donor assignment while achieving fast runtimes (Fig. 3B, Table 6) and low memory (Fig. 3C, Table 6) usage even when incorporating variants with coverage as low as 10 reads.

**Figure 3.**
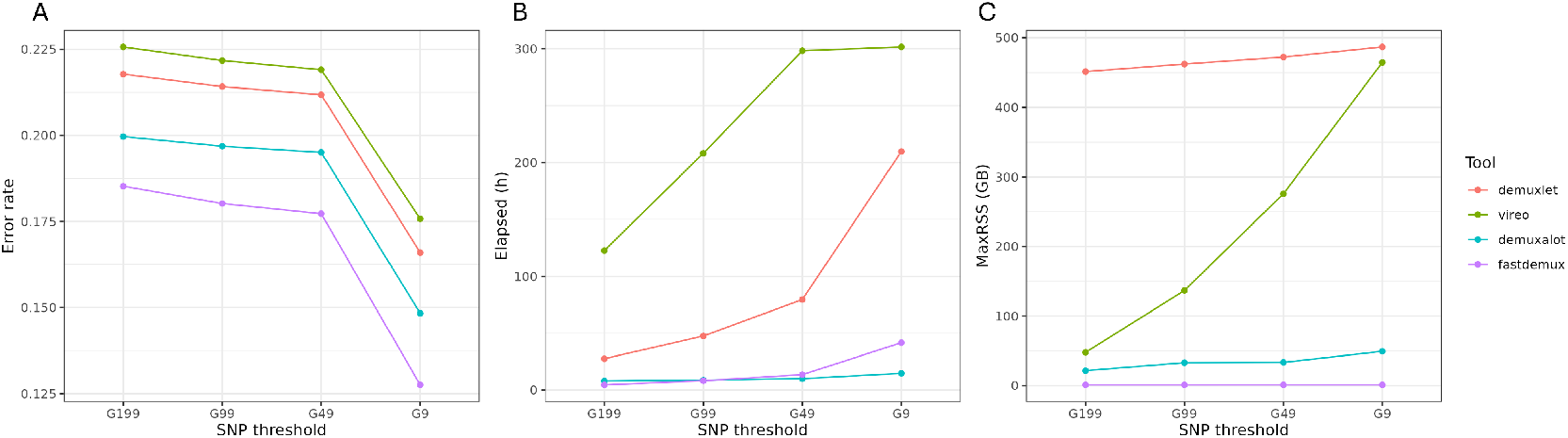
Demultiplexing performance as a function of genotype SNP density. x-axis represents the progressively relaxed SNP coverage thresholds (G199, G99, G49, and G9) using a higher number of SNPs, and the y-axis compares across demultiplexing tools: (A) Total-droplet error rate as a function of genotype SNP filtering thresholds, evaluated relative to each tool’s baseline run using the G9-filtered genotype set. (B) Total elapsed runtime in hours. (C) Peak resident memory usage in gigabytes (MaxRSS). All tools were evaluated on the same pooled 8-donor scRNA-seq dataset.

The high computational efficiency of `fastdemux` using a large number of lower-coverage SNPs can be leveraged for multiplexing scATAC-seq data, in which genetic variants are typically observed at substantially lower coverage than in scRNA-seq. We evaluated `fastdemux` on multiplexed scATAC-seq data from a PBMC library comprising 12 individuals [3]. Using an overloaded library with a mean of 64,908 fragments per cell, `fastdemux` successfully assigned 17,869 singlets to donors with a total user time of 31.33 hours and a maximum memory usage of 1.26 GB. We attempted to demultiplex this library using demuxalot, but this tool requires an NH field tag on the alignment file that is not present in the scATAC-seq data. We also note that for a library of this size and coverage, utilizing demuxlet and vireo is not computationally practical.

## Discussion

We developed `fastdemux,` a scalable genotype-based demultiplexing method designed to address the growing computational challenges of pooled single-cell genomics experiments. Using a diagonal linear discriminant analysis (DLDA) framework, `fastdemux` efficiently assigns droplets to their most likely donor of origin while supporting explicit modeling of singlets, doublets, and higher-order multiplets within a unified statistical formulation. Across a well-characterized pooled scRNA-seq dataset, `fastdemux` achieved demultiplexing accuracy comparable to or exceeding that of the most commonly used tool, demuxlet [11], while substantially reducing computational runtime and total memory usage.

A central contribution of `fastdemux` is its computational efficiency and streaming algorithm. In contrast to likelihood-based demultiplexing methods such as demuxlet, which require repeated evaluation of per-cell likelihoods across donors and genotype configurations [11], the DLDA formulation enables `fastdemux` to operate through linear scoring functions that aggregate allelic evidence across variants while reading the input files. By exploiting the conditional independence assumptions inherent to DLDA and operating on aggregated, standardized allele count summaries rather than read-level likelihoods,`fastdemux` avoids expensive iterative probability evaluations and substantially reduces computational overhead. These properties translate into favorable scaling behavior as the number of cells, donors, and informative variants increases, making `fastdemux` particularly well-suited for large-scale population based single-cell studies.

Importantly, improvements in computational performance were not achieved at the expense of demultiplexing accuracy. Across all benchmarked tools, `fastdemux` recovered a comparable or greater number of intended singlets and exhibited the lowest fraction of unintended droplets, with consistent donor-level recovery across individuals. Concordance with expression-derived sex annotations further indicated that `fastdemux` does not introduce systematic assignment biases relative to existing methods, and observed sex misclassification rates were driven primarily by limitations intrinsic to expression-based sex inference rather than demultiplexing performance. Together, these results demonstrate that `fastdemux` preserves biologically meaningful donor assignments while offering substantial gains in computational efficiency.

Our analyses further highlight how demultiplexing performance depends on sequencing depth and genotype information, consistent with the statistical foundations of genotype-based assignment. All methods exhibited reduced accuracy under extreme read downsampling, reflecting limited allelic information available per droplet at very low coverage. However,`fastdemux` consistently achieved lower error rates than other tools at moderate to high sequencing depths, indicating that its discriminant framework effectively integrates available allele-level evidence when suffcient signal is present. Similarly, while all tools showed increased error rates as genotype SNP density decreased, `fastdemux` demonstrated greater robustness under stringent SNP filtering. Importantly, we highlight that`fastdemux` can efficiently perform accurate donor assignment using a high number of low-coverage SNPs, which can also be utilized for demultiplexing scATAC-seq data with naturally sparse and low coverage genetic variations.

A key methodological advantage of `fastdemux` is its natural extension to multiplet detection. By expressing doublet and higher-order multiplet discriminant scores directly in terms of singlet scores,`fastdemux` can evaluate arbitrary M-let configurations within the same DLDA framework without introducing combinatorial complexity. This formulation allows`fastdemux` to detect not only doublets but also higher-order multiplets in a principled and computationally tractable manner, an increasingly important capability as cell loading densities and experimental throughput continue to increase. This contrasts with approaches that focus primarily on singlet–doublet discrimination and aligns with the growing recognition that multiplets represent a non-negligible fraction of droplets in large-scale experiments [33].

Despite these strengths, several limitations should be considered. Like all genotype-based demultiplexing approaches, `fastdemux` relies on the availability of informative genetic variation between donors. Related individuals pose an inherent challenge for genotype-based demultiplexing, as shared haplotypes reduce the discriminative power of SNP-based assignment regardless of the underlying statistical framework. These limitations are not unique to `fastdemux` but reflect fundamental constraints of demultiplexing methods based on natural genetic variation. Nevertheless,`fastdemux` performs well when a large number of low coverage SNPs are available which is a common situation in scATAC-seq.

In summary, `fastdemux` provides a computationally efficient and statistically principled alternative to existing demultiplexing tools, enabling accurate donor and multiplet assignment while substantially reducing runtime and memory requirements. By addressing key scalability limitations in the commonly used tools, `fastdemux` facilitates the application of genetic demultiplexing to increasingly large pooled single-cell datasets, helping to expand the practical scope of population-level multiplexed single-cell experimental designs.

## Acknowledgments

We thank members of the Pique-Regi/Luca group for helpful comments and discussions, and for providing feedback on the method performance. This work was funded by R01GM109215 and R01HL162574 (FL, RPR), and CZI Ancestry Networks (FL, RPR). AI tools were used to edit portions of the initial draft of the manuscript.

## Code Availability

The method is available at: https://github.com/piquelab/fastdemux, and the code for the benchmarks and comparisons can be found at: https://github.com/piquelab/fastdemux_bench

## Notes

https://github.com/piquelab/fastdemux

### Competing Interest Statement

The authors have declared no competing interest.

