## Supplement for "fastdemux: Robust SNP-based demultiplexing of single-cell population genomics data"

### Supplements

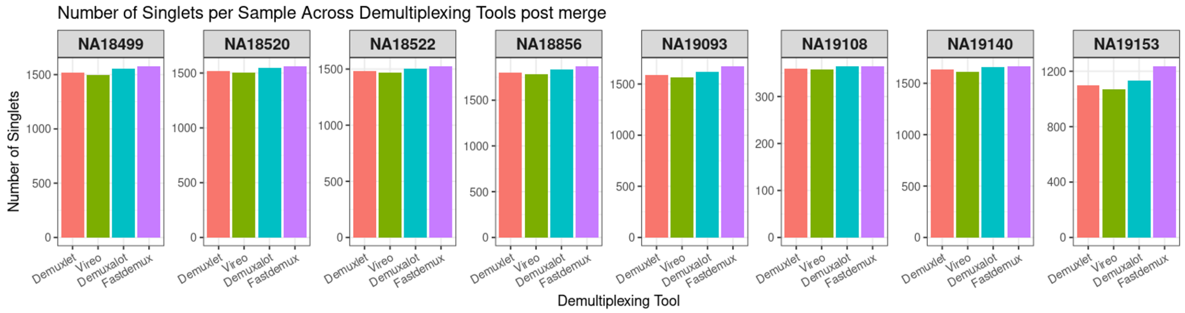


Figure S1: **Donor-level recovery of singlets across demultiplexing tools.** Number of singlets recovered per donor across demuxlet, vireo, demuxalot, and `fastdemux` after merging demultiplexing outputs.

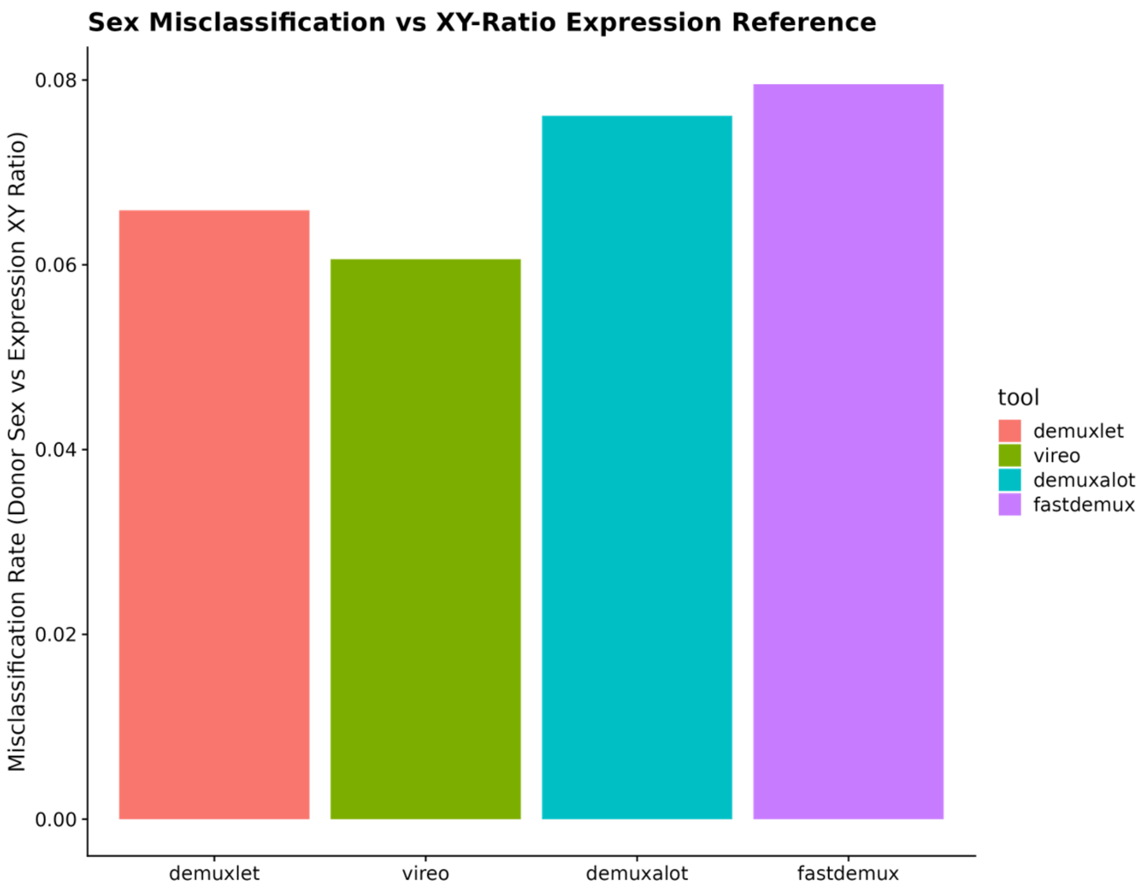


Figure S2: **Expression-based sex assignment quality control.** Concordance between donor sex metadata and expression-based sex inference across demultiplexing tools. Sex was inferred independently of demultiplexing by computing per-cell X/Y expression ratios from pooled scRNA-seq data, followed by log transformation and unsupervised k-means clustering to define a global sex assignment threshold.

Table 1: **Computational resource usage across demultiplexing tools.** Comparison of demultiplexing tools showing total elapsed runtime in hours (Elapsed), cumulative CPU time used across all cores (TotalCPU), and peak resident memory usage in gigabytes (MaxRSS), evaluated under comparable computational settings.

| Tool | Elapsed (h) | TotalCPU (h) | MaxRSS (GB) |
|---|---|---|---|
| demuxlet | 209.66 | 207.03 | 487.03 |
| vireo | 301.55 | 299.77 | 464.70 |
| demuxalot | 14.78 | 11.36 | 49.33 |
| `fastdemux` | 41.68 | 46.64 | 1.02 |

Table 2: **Summary of droplet assignment categories across demultiplexing tools.** Counts of droplets assigned to each category by demuxlet, vireo, demuxalot, and `fastdemux`. Reported categories include ambiguous droplet calls (AMB), doublets (DBL), unintended droplets (droplets assigned to samples outside the intended pool), intended singlets (Intended SNG), and corresponding counts after merging Seurat count data with demultiplexed results.

| Tool | AMB | DBL | Unintended droplets | Intended SNG | Unintended merged droplets | Merged SNG | Intended merged SNG |
|---|---|---|---|---|---|---|---|
| demuxlet | 1711663 | 67797 | 3928 | 22145 | 2186 | 10987 | 10987 |
| vireo | 800 | 2212 | 3012 | 10861 | 2315 | 13173 | 10858 |
| demuxalot | 0 | 2615 | 0 | 11258 | 1954 | 11219 | 11219 |
| `fastdemux` | 28511 | 20046 | 896 | 27678 | 1683 | 11493 | 11490 |

Table 3: **Fraction of intended and unintended droplet assignments across demultiplexing tools.** Proportion of droplets assigned as intended singlets and unintended droplets after post-merge processing.

| Tool | Intended merged SNG | Unintended merged droplets |
|---|---|---|
| demuxlet | 0.83 | 0.17 |
| vireo | 0.82 | 0.18 |
| demuxalot | 0.85 | 0.15 |
| `fastdemux` | 0.87 | 0.13 |

Table 4: **Number of singlets recovered per donor across demultiplexing tools.**

| Sample ID | Demuxlet | Vireo | Demuxalot | `fastdemux` |
|---|---|---|---|---|
| NA18499 | 1515 | 1495 | 1557 | 1579 |
| NA18520 | 1518 | 1505 | 1544 | 1564 |
| NA18522 | 1484 | 1469 | 1507 | 1527 |
| NA18856 | 1797 | 1784 | 1838 | 1873 |
| NA19093 | 1586 | 1566 | 1619 | 1676 |
| NA19108 | 360 | 358 | 364 | 365 |
| NA19140 | 1631 | 1611 | 1658 | 1668 |
| NA19153 | 1096 | 1070 | 1132 | 1238 |

Table 5: **Computational resource usage across demultiplexing tools at varying sequencing depths.** Resource utilization for demuxlet, vireo, demuxalot, and `fastdemux` evaluated across downsampled read fractions. Reported metrics include mean reads per cell, total elapsed runtime in hours (Elapsed), cumulative CPU time used across all cores (TotalCPU), and peak resident memory usage in gigabytes (MaxRSS), evaluated under comparable computational settings.

| Tool | Read Fraction | Reads per cell | Elapsed (h) | TotalCPU (h) | MaxRSS (GB) |
|---|---|---|---|---|---|
| demuxlet | 50% | 38,119 | 99.144 | 98.530 | 261.961 |
| demuxlet | 30% | 22,871 | 55.924 | 55.582 | 164.113 |
| demuxlet | 10% | 7,624 | 15.630 | 15.488 | 58.196 |
| demuxlet | 5% | 3,812 | 6.879 | 6.798 | 47.660 |
| demuxlet | 1% | 762 | 0.988 | 0.939 | 42.481 |
| vireo | 50% | 38,119 | 165.935 | 165.178 | 337.991 |
| vireo | 30% | 22,871 | 106.018 | 105.454 | 245.078 |
| vireo | 10% | 7,624 | 24.563 | 24.466 | 90.864 |
| vireo | 5% | 3,812 | 8.942 | 8.888 | 46.530 |
| vireo | 1% | 762 | 2.971 | 2.933 | 27.213 |
| demuxalot | 50% | 38,119 | 7.245 | 7.147 | 35.098 |
| demuxalot | 30% | 22,871 | 5.526 | 5.463 | 30.618 |
| demuxalot | 10% | 7,624 | 2.427 | 2.399 | 25.038 |
| demuxalot | 5% | 3,812 | 1.688 | 1.666 | 23.512 |
| demuxalot | 1% | 762 | 0.985 | 0.942 | 22.028 |
| `fastdemux` | 50% | 38,119 | 16.112 | 27.766 | 0.903 |
| `fastdemux` | 30% | 22,871 | 10.386 | 18.292 | 0.844 |
| `fastdemux` | 10% | 7,624 | 5.043 | 7.957 | 0.763 |
| `fastdemux` | 5% | 3,812 | 3.254 | 4.848 | 0.734 |
| `fastdemux` | 1% | 762 | 0.642 | 1.105 | 0.710 |

Table 6: **Computational resource usage across VCF SNP coverage thresholds.** Runtime and memory utilization for demuxlet, vireo, demuxalot, and `fastdemux` evaluated using genotype VCF files filtered at progressively relaxed SNP coverage thresholds (G199, G99, G49) and the default threshold (G9). Reported metrics include the number of retained SNPs (No. SNPs), total elapsed runtime in hours (Elapsed), cumulative CPU time used across all cores (TotalCPU), and peak resident memory usage in gigabytes (MaxRSS), evaluated under comparable computational settings.

| Tool | SNP threshold | No. SNPs | Elapsed (h) | TotalCPU (h) | MaxRSS (GB) |
|---|---|---|---|---|---|
| demuxlet | G199 | 334,896 | 27.63 | 27.32 | 451.45 |
| demuxlet | G99 | 745,352 | 47.60 | 47.22 | 462.31 |
| demuxlet | G49 | 1,470,358 | 79.59 | 79.32 | 472.44 |
| demuxlet | G9 | 4,694,878 | 209.66 | 207.03 | 487.03 |
| vireo | G199 | 334,896 | 122.59 | 122.06 | 47.77 |
| vireo | G99 | 745,352 | 208.11 | 207.26 | 136.82 |
| vireo | G49 | 1,470,358 | 298.18 | 296.25 | 275.82 |
| vireo | G9 | 4,694,878 | 301.55 | 299.77 | 464.70 |
| demuxalot | G199 | 334,896 | 8.00 | 7.87 | 21.57 |
| demuxalot | G99 | 745,352 | 8.79 | 8.62 | 33.00 |
| demuxalot | G49 | 1,470,358 | 10.08 | 9.77 | 33.45 |
| demuxalot | G9 | 4,694,878 | 14.78 | 11.36 | 49.33 |
| `fastdemux` | G199 | 334,896 | 4.58 | 6.12 | 0.99 |
| `fastdemux` | G99 | 745,352 | 8.26 | 11.58 | 0.99 |
| `fastdemux` | G49 | 1,470,358 | 13.62 | 19.98 | 1.03 |
| `fastdemux` | G9 | 4,694,878 | 41.68 | 46.64 | 1.02 |
